# Evaluation of the taxonomic accuracy and pathogenicity prediction power of 16 primer sets amplifying single copy marker genes in the *Pseudomonas syringae* species complex

**DOI:** 10.1101/2022.11.01.514712

**Authors:** Chad Fautt, Kevin L. Hockett, Estelle Couradeau

**Affiliations:** Department of Plant Pathology and Environmental Microbiology, Pennsylvania State University, University Park, Pennsylvania, United States; Department of Ecosystem Science and Management, Pennsylvania State University, University Park, Pennsylvania, United States; Intercollege Graduate Degree Program in Ecology, Pennsylvania State University, University Park, Pennsylvania, United States

## Abstract

The Pseudomonas syringae species complex is comprised of several closely related species of bacterial plant pathogens. Here, we use in-silico methods to assess 16 PCR primer sets designed for broad identification of isolates throughout the species complex. We evaluate their in-silico amplification rate in 2,161 publicly available genomes, the correlation between pairwise amplicon sequence distance and whole genome average nucleotide identity (ANI), and we train naïve Bayes classification models to quantify classification resolution. Further, we show the potential for using single amplicon sequence data to predict an important determinant of host specificity and range, type III effector protein repertoires.

## INTRODUCTION

The *Pseudomonas syringae* species complex (PSSC) consists of many closely related plant pathogens (Sarkar and Guttman 2004). With host ranges and symptomology that can overlap, proper identification of isolates can be difficult (Morris et al. 2019). Aside from whole genome sequencing, which is costly and often impractical for routine identification, marker gene sequencing is the most effective method for specific and sub-specific classification of unknown PSSC isolates (Berge et al. 2014; Borschinger et al. 2016; Guilbaud et al. 2016). This method has been used to aid in the identification of new pathogenic species and expand our view of host range in known pathogens (Dutta et al. 2018; Keshtkar, Khodakaramian, and Rouhrazi 2016; Moretti, Fakhr, and Buonaurio 2012), highlighting the importance of amplicon sequencing for broadening our understanding of the species complex. However, although there have been many proposed PCR primer sets designed to amplify broadly within the species complex, there are open questions about the relative performance of each. Specifically, it’s not clear if all primers allow for reliable amplification for all phylogroups in the species complex, or which primer sets allow for the greatest classification resolution. Further, while a primary goal of pathogen identification is often to predict pathogenic potential of the unknown isolate, it isn’t known if the classification resolution obtained by any of the currently published primers is sufficient to predict genomic features associated with host range and virulence.

While most of the primer sets evaluated in this study were originally designed for use with multi-locus sequence typing (MLST) (Sarkar and Guttman 2004; Yan et al. 2008), there has been continuous interest in classifying isolates with a single marker gene. In this regard, recombination rates and phylogenetic congruence have been used as metrics to suggest that genes such as citrate synthase (CTS) (Berge et al. 2014) and RNA polymerase sigma factor rpoD (Parkinson et al. 2011) can be used by themselves to accurately place unknown PSSC isolates into phylogroups. An in-depth comparison of these primer sets as tools for classification has not been conducted, however, leaving it an open question as to which performs best.

Often, an implicit goal of bacterial isolate identification is to gain some insight into its functionality or ecological significance for the environment it was isolated from. In this vein, assuming a phylogenetic placement with sufficient resolution, prediction of an isolate’s gene content can be made, providing insight into functional capacity. This concept has been demonstrated with Picrust2, in which improved functional predictions were achieved over Picrust1 solely from increased resolution in genome prediction (Douglas et al. 2020). As PCR primer sets designed specifically for PSSC are used because they offer even greater phylogenetic resolution over those targeting 16S rRNA genes (the marker gene used by Picrust2), we hypothesized that specific genes known to affect host range and virulence could be predicted in genomes based solely on amplicon sequences derived from commonly used PCR primer sets.

In PSSC, pathogenicity is determined in large part by the type III effector proteins (T3Es) carried by the pathogen (Hulin et al. 2018) and therefore predicting T3E repertoires could provide valuable information about host range and specialization of an unknown isolate. While many pathogens in the species complex carry 30-40 T3Es, only effectors AvrE, HopM, and HopAA are considered part of the core PSSC genome of PSSC and are thought to confer general virulence to plants (Dillon et al. 2019). Presence/absence of the other T3Es play a role in host adaptation and are more variable in the species complex, suggesting that if there is a taxonomic signature associated to their presence, an infraspecific classification is needed to accurately predict it. It is currently not known what phylogenetic resolution is needed to accurately predict T3E repertoires, or whether any published primer sets might allow high enough resolution to meet this threshold.

Here, we perform in-silico tests to assess the performance of 16 previously published PCR primer sets, targeting eight marker genes, against 2,161 PSSC genomes. Metrics used for assessment of phylogenetic classification were amplification rate, congruence of pairwise amplicon distance with average nucleotide identity (ANI), and performance of naïve Bayes classifiers trained on in-silico amplicon data. We also investigate the potential for functional prediction from amplicon data by analyzing Jaccard similarity of T3E repertoires at the level of phylogenetic resolution achieved by each classifier and show that isolates not included in the training dataset can be accurately placed above phylogroup level and prediction of both T3E repertoire size and content is often possible, with presence/absence of 77 T3E subfamilies being correctly predicted 75-100% of the time among genomes in our 2,161 genome training set, and 79.2-94.8% of the time among 5 genomes, representing phylogroups 2,3 and 4, outside our training set.

Overall, we find that some published primer sets have substantial blind spots in the lineages they can amplify. However, many primers tested can both amplify broadly throughout the species complex and be used to classify isolates beyond the phylogroup level, allowing accurate prediction of the T3Es carried by unknown PSSC isolates. Our results suggest that for highest classification resolution throughout the species complex, resulting in the most consistent T3E repertoire prediction accuracy, primer sets targeting the genes gapA, gyrB (Hwang et al. 2005) and PGI (Yan et al. 2008) should be considered as the optimal primer sets.

## EXPERIMENTAL PROCEDURES

2,467 Genomes labeled as belonging to the *Pseudomonas syringae* group were obtained from the RefSeq database from NCBI in November 2021. Genomes were checked for completeness and assembly quality with BUSCO v 5.3.2 using default settings and the pseudomonadales_odb10 lineage. Genomes scoring > 99 made up the final dataset used for assessing primers. As the majority of genomes used were not assigned to a phylogroup, phylogroups were assigned based on average nucleotide identity (ANI) with phylogroup reference genomes produced by (Berge et al. 2014). While Berge et al. suggest taking a simple nearest neighbor approach to assigning phylogroup, 173 genomes within our dataset shared less than 95% ANI with any phylogroup reference genome, indicating that they were either misclassified at the time of depositing into GenBank as belonging to PSSC, or that they might represent new phylogroups. As a result, these genomes were left unassigned to a phylogroup. Eventually a curated set of 2,161 genomes was used, with 1,988 assigned to a phylogroup (supplementary table 1 contains the accession numbers of these genomes and assigned phylogroups, along with ANI clusters and T3E gene content described below).

**Table 1.**
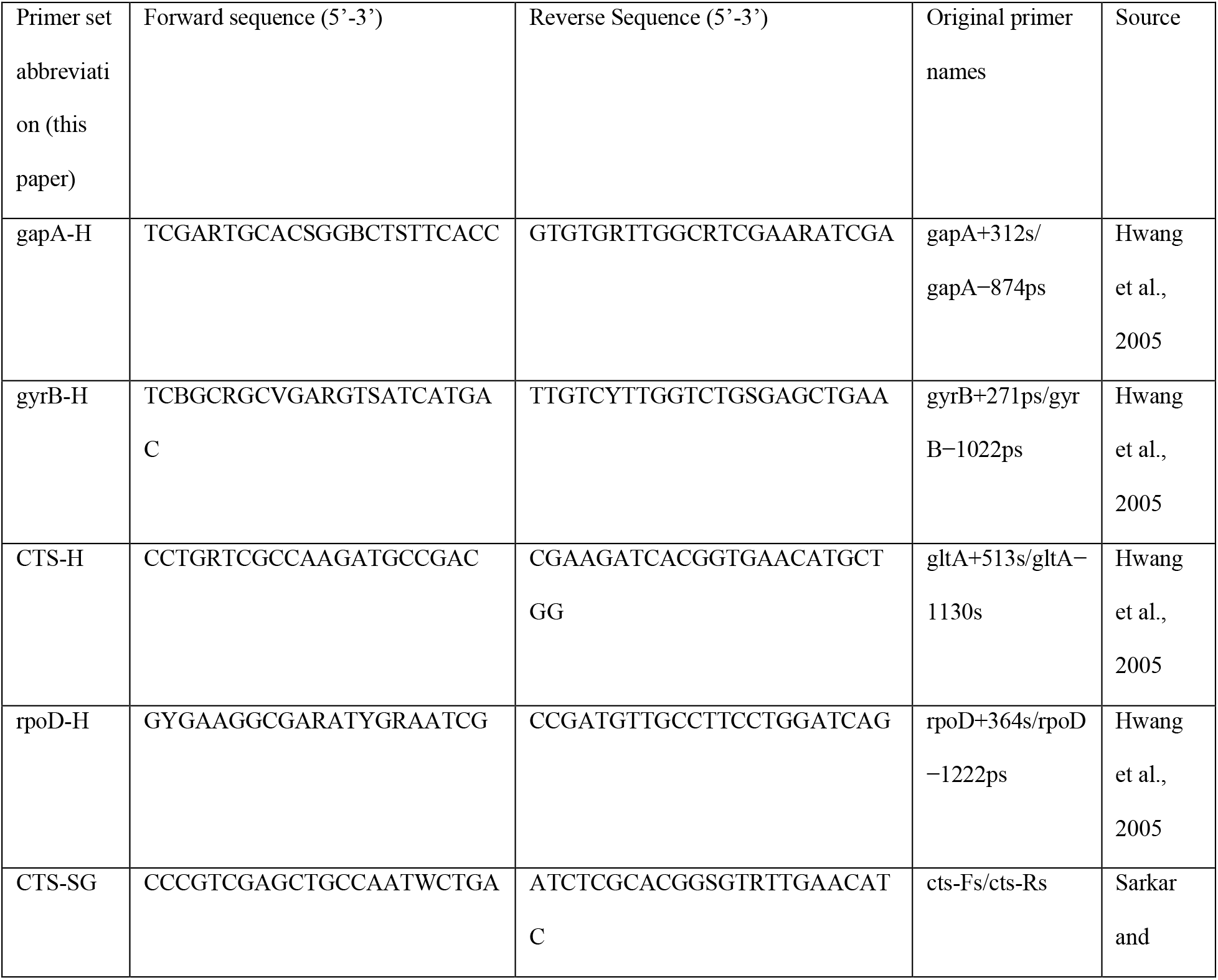
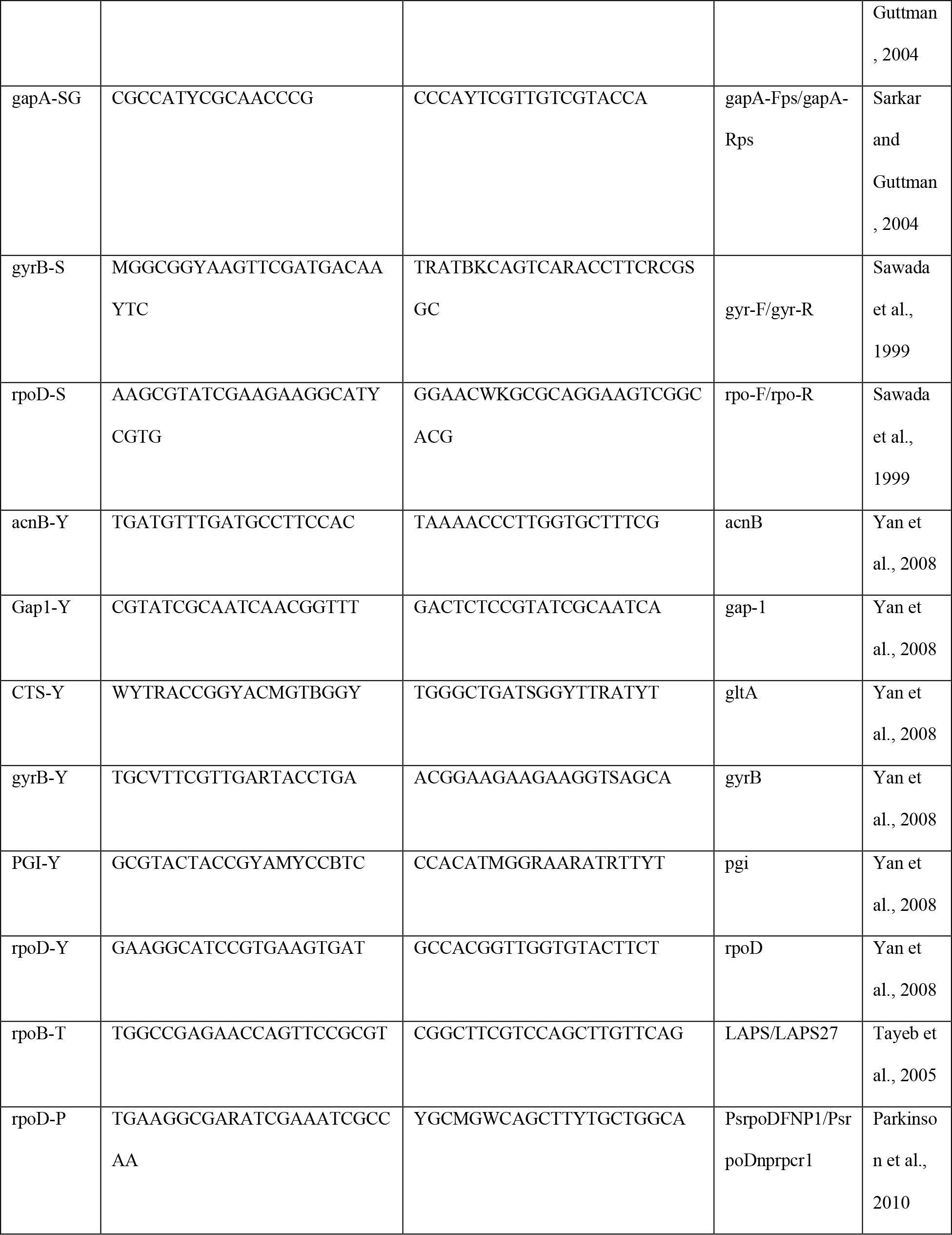
Primer sets used in this study

In-silico PCR was performed with *in_silico_PCR* (Ozer 2020) allowing for one mismatch per primer. The identity of amplicons was confirmed by visually inspecting multiple sequence alignments performed with MAFFT, using Geneious 2019.1.3 (https://www.geneious.com). The amplification rate reported is the percentage of the 2,161 genomes that resulted in successful amplification of the target gene fragment. Primers included in this study are given in table 1.

Pairwise ANI values for all genomes were computed with fastANI v1.33 (Jain et al. 2018). For each primer set with amplification rate greater than 50%, amplicon sequences were aligned using MAFFT version 7 (Katoh and Standley 2013) with the options ‘globalpair’ and a ‘maxiterate’ of 1000. For amplicon sequence similarity, pairwise hamming distances for the aligned sequences were then computed with the ‘DistanceMatrix’ function in the R package DECIPHER (Wright 2015). To quantify the correlation between amplicon sequence distances and ANI of the source genomes, the mean squared deviation of amplicon sequence similarity from ANI was computed as the sum of squared distances between the two values for each genome pair. As ‘DistanceMatrix’ reports distance in the range of 0-1, ANI values were normalized to the same range by dividing by 100.

Using QIIME2’s v2022.2 feature-classifier (Bolyen et al. 2019), naïve Bayes classifiers were trained on the unaligned *in-silico* amplicon sequences generated above. Primer sequences were left untrimmed from amplicons. Classification models require taxonomic descriptions of known genomes for training, however, nomenclature in PSSC is largely inconsistent (Gomila et al. 2017), which can significantly reduce the predictive power of classification models. We therefore implemented instead a strict hierarchical taxonomy based on ANI, generated by the hierarchical the clustering algorithm utilized by LINbase (Tian et al. 2020). Briefly, a randomly selected genome in the set is assigned to clusters representing twenty ANI values from 80-99%, each given a numeric signature of ‘0’. All other genomes are iteratively assigned to clusters based on the closest already-assigned genome, and given the same numeric signature for all ANI values up to the point which the two genomes differ, wherein the numeric signature is iterated (e.g. if genome #2 shares 97.5% ANI with genome #1, genome #2 will be assigned to cluster ‘0’ along with genome #1 for all ANI values except 98%, where it will be assigned a numeric signature of ‘1’). The resulting taxonomy file consisting of twenty taxonomic levels representing ANI values from 80-99% (see supplementary table 1).

Reference sequences for T3E subfamilies included in the *Pseudomonas syringae* type III effector compendium (PsyTEC) (Laflamme et al. 2020) were aligned using MAFFT with default settings, and the alignments input into the HMMER v3.1b2 (hmmer.org) function HMMbuild to generate HMM profiles. Using HMMsearch, these 77 HMMs were run on the set of 2,161 genomes and an e-value of 10^−20^ was used as the threshold for considering a subfamily to be present in a genome.

## RESULTS AND DISCUSSION

As all primers used in this study were designed to work broadly on strains within the species complex, we first tested amplification rate of each primer set. Surprisingly, when tested on a comprehensive set of genomes representing the full known diversity of PSSC, 7/16 primer sets tested had an amplification rate of less than 50% (Fig 1a). These primers were removed from any further tests. For the remaining 9 primers, performance was substantially better, with amplification rates ranging from 91.37% (rpoD-P) to 100% (rpoD-H). These large differences in amplification rate are likely due to the significant degeneracy built into the best performing primer sets (table 1 and supplementary table 2) and highlights the importance of considering the known diversity of PSSC when designing primers.

**Table 2.** Summary of classification test results

| Primer set | Mean ANI resolution | SD |
| --- | --- | --- |
| gyrB-H | 97.5970987 | 0.59560943 |
| CTS-SG | 97.2226044 | 0.59653389 |
| CTS-Y | 97.2277597 | 0.65161671 |
| gapA-H | 97.5547818 | 0.5947916 |
| rpoD-P | 97.6722441 | 0.66541184 |
| PGI-Y | 97.9319664 | 0.43925883 |
| gyrB-Y | 97.5712166 | 0.57012883 |
| rpoD-H | 97.6307265 | 0.69616513 |
| CTS-H | 97.2234903 | 0.80563859 |

**Figure 1.**
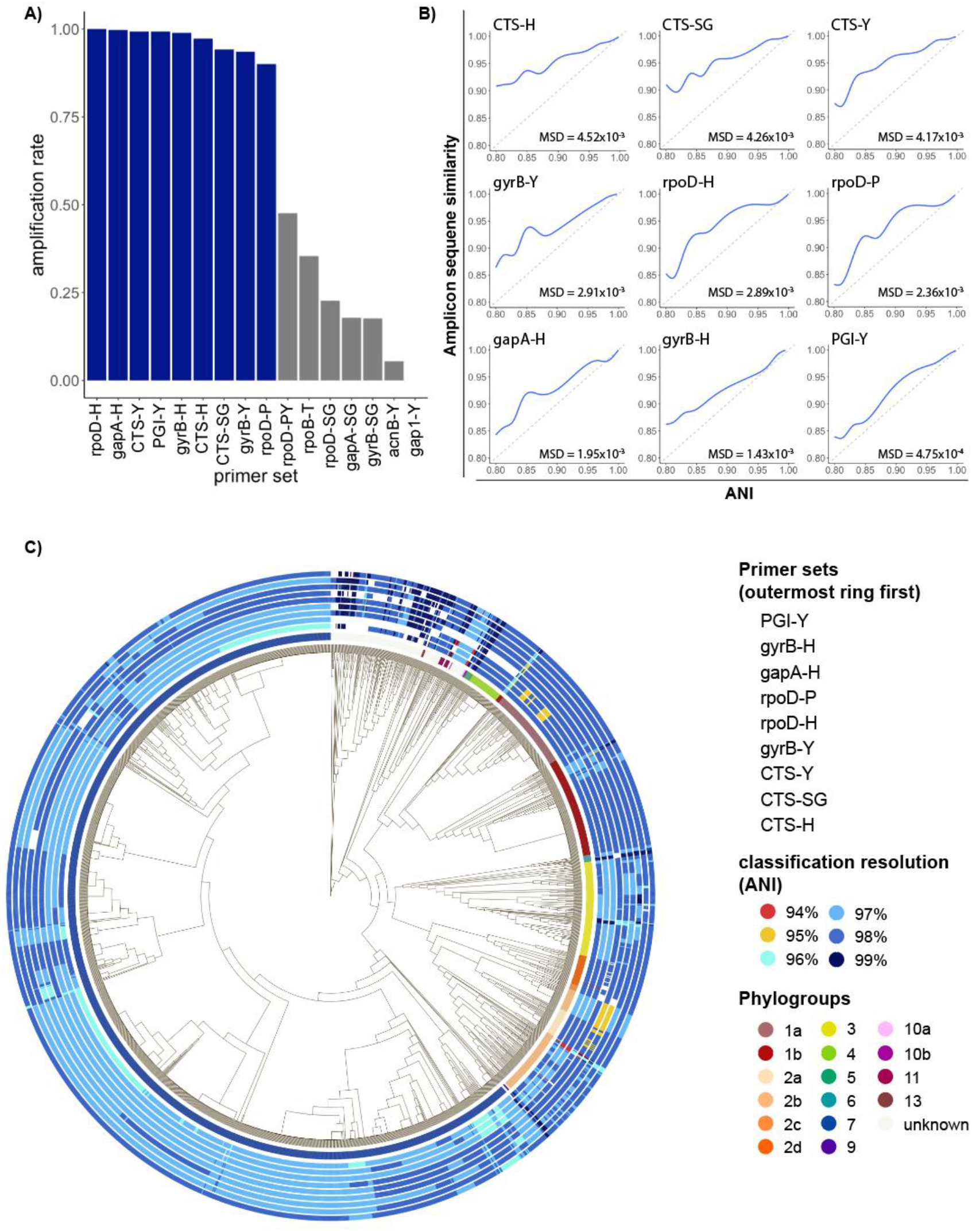
Results of amplicon-based classification tests. **a)** proportion of genomes with successful amplification, allowing 1 mismatch per primer. Primer sets producing amplicons in more than 90% of genomes are highlighted in blue. Primer sets in grey were omitted from further analysis. **b**) Generalized additive models summarizing relationship between pairwise amplicon similarity and whole genome ANI. Mean squared deviation (MSD) of amplicon similarity from ANI is shown in the lower left corner, dashed grey line represents MSD = 0. **c)** Core genome phylogeny for all genomes used in this study. Innermost ring annotates phylogroups, while each outer ring represents the classification resolution obtained when amplicon sequences from each genome were identified with a Bayes classifier. White rectangles represent genomes from which in-silico amplification was unsuccessful.

When choosing a marker gene, or region of a marker gene, to use for identification purposes, an important consideration is the level of conservation found within the region, as regions that are too conserved result in reduced taxonomic resolution. To investigate the amount of conservation found within amplicons generated by each primer set, we compared pairwise amplicon similarity (represented by their Jaccard index) with ANI of the genomes from which the amplicons were derived (Fig 1b). Of the primers tested, amplicons from citrate synthase showed the highest level of conservation, indicating they might not provide the best resolution when used for classification contrary to previous suggestions (Berge et al., 2014). On the other hand, amplicons generated by PGI-Y, targeting phosphoglucose isomerase, exhibited a mean squared deviation from ANI almost 10 times lower than any primer targeting CTS (Fig 1b), indicating a very good congruence between the diversity at this locus and the one retrieved at the genome level.

To compare performance of primers in classifying individual strains throughout PSSC, classification models were trained on amplicons generated by each primer set, and then used to classify each strain in the training set. The relative performance of the classification models mirrored the amount of conservation observed above (fig 1b), although the practical differences in classification resolution were minimal (table 2). Remarkably, every primer set allowed for classification beyond phylogroup level, with mean ANI of predicted clades ranging from 97.22% (CTS-H, CTS-Y & CTS-SG) to 97.93% (PGI-Y). Surprisingly, while the CTS gene has been suggested to be a particularly informative marker gene for PSSC (Berge et al., 2014), the three primers targeting this gene performed slightly below the other primers tested. While mean performance of classification models suggest PGI-Y as the best primer set, it doesn’t consider biases in the representation of each phylogroup in our dataset, and so we sought to analyze any discrepancies in primer performance among phylogroups (Fig 1c).

For strains belonging to Phylogroup 1, all primers performed well, amplifying every strain tested and classifying most to 98% ANI (Fig 1c). An exception to the strong performance can be seen for 2 subclades which CTS-Y and CTS-SG were only able to classify at 95%. Overall, the best performing primers for phylogroup 1 were gapA-H and gyrB-H.

In Phylogroup 2, performance was more variable. Both rpoD-P and rpoD-H were unable to classify strains in a well sampled clade of phylogroup 2b above 95% ANI, and gyrB-Y was unable to amplify several strains within phylogroup 2d. As seen in phylogroup 1, the best performing primers for phylogroup 2 were gapA-H and gyrB-H.

Phylogroup 3 strains were successfully amplified by every primer, with the exception of four strains that gyrB-H was unable to amplify from. Overall, PGI-Y and gyrB-Y were the best performing primers for strains in phylogroup 3.

All primers performed equally well for phylogroup 4 strains, classifying to 98% ANI. gyrB-Y, however, failed to amplify from every strain in this phylogroup.

Our dataset contained only nine phylogroup 5 strains, but every primer set was able to amplify and classify each one to 98-99% ANI.

As with phylogroup 4, gyrB-Y was the only primer set unable to amplify and classify to 97-99% every strain in phylogroup 6.

Within phylogroup 7, PGI-Y performed the best, classifying 89% (1,189/1,331) of strains to 98% ANI, while all other primers generally classified strains in this phylogroup to 97%.

Phylogroups 9, 10, 11, and 13 were underrepresented in the dataset and so conclusions are difficult to draw about primer performance. Additionally, there were several strains not assigned to a phylogroup in our dataset. Among these strains, rpoD-H and gapA-H exhibited the highest amplification rates, and classification resolution ranging from 97-99% ANI.

No single primer set universally outperformed the rest, and as such the suspected identity of an unknown isolate should be considered when choosing the appropriate set of primers to use for classification. However, PGI-Y, gapA-H, and gyrB-H generally performed well throughout the species complex and should be considered the default choices for highest classification resolution.

As classification based on single amplicon sequences resulted in fairly high genomic resolution (97-98% ANI), we sought to explore the possibility of predicting gene content of a target strain using gene content of predicted relatives. As type III effector proteins (T3Es) are important determinants of virulence in PSSC (Dillon et al. 2019; Lindeberg, Cunnac, and Collmer 2009), we focused here on T3Es by first assessing the prevalence of each T3E subfamily within genomic clusters sharing at least 98% ANI (fig 2). While there were several clusters that contained only a single genome (i.e., no genome in the dataset shared more than 98% ANI with them), even among better represented clusters, there was considerable similarity in T3E repertoires. This suggested that unknown isolates placed into these clusters should exhibit a predictable T3E repertoire. Perhaps not surprisingly, clusters representing phylogroups that contain most of the agricultural pathogens within PSSC exhibit many more T3Es as well as a greater diversity in repertoires between strains, indicated by the average Jaccard index within a given cluster (fig 2). When the T3E repertoires of the 2,161 genomes were compared against the consensus repertoires (defined by taking the most common state of each T3E subfamily; absent or present) of their 98% ANI clusters, 75.3-100% of actual T3E states recapitulated the intra-cluster consensus (fig 3).

**Figure 2.**
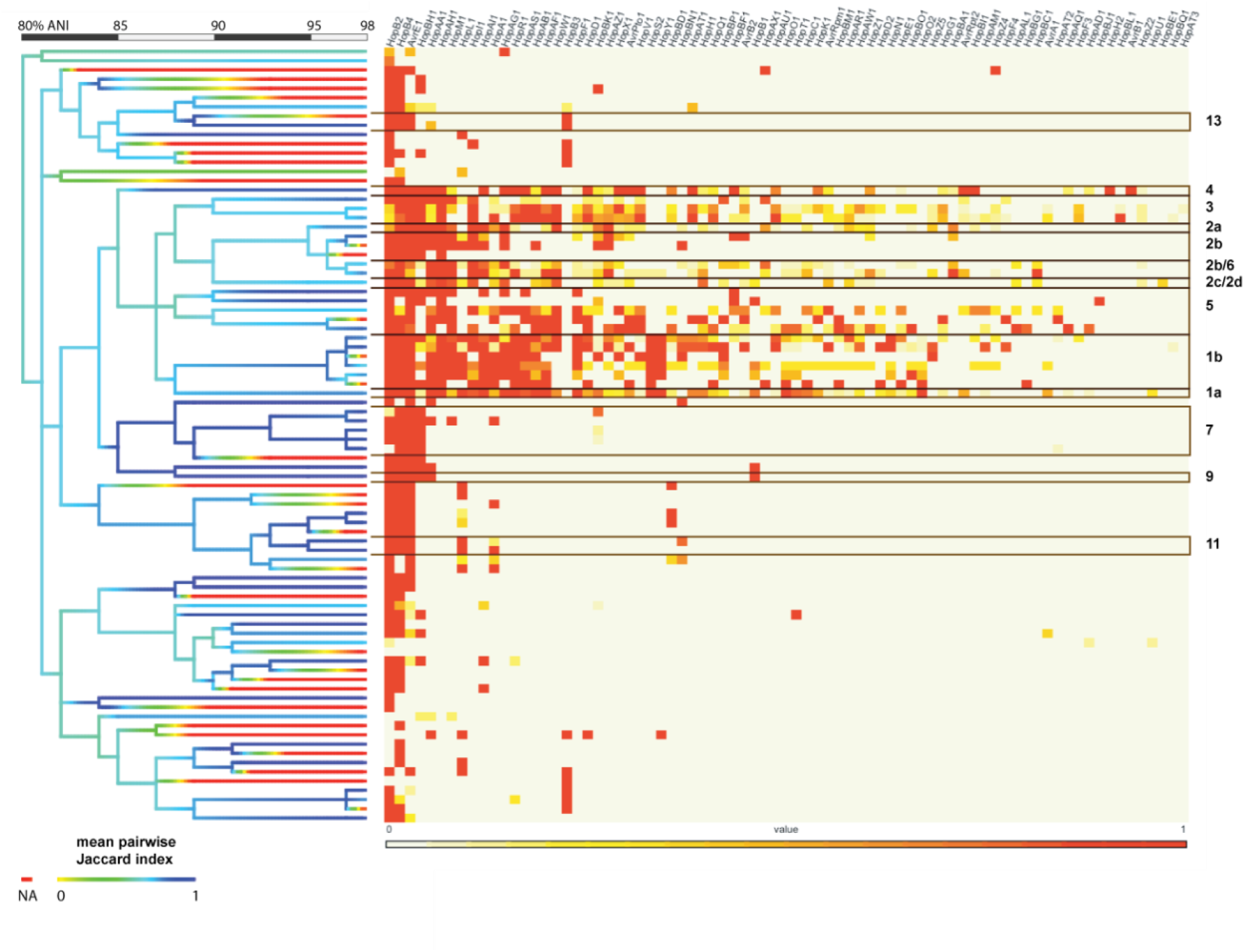
Distribution of type III effector proteins throughout PSSC. Heatmap shows frequency of each T3E subfamily, with each row as a cluster of genomes sharing 98% ANI. Black outlines indicate phylogroups found in each row. Cladogram on left represents ANI-based clusters of genomes from 80-98% ANI and is colored by mean Jaccard index of genome pairs within each branch. Red branches indicate singletons, for which Jaccard index could not be calculated.

**Figure 3.**
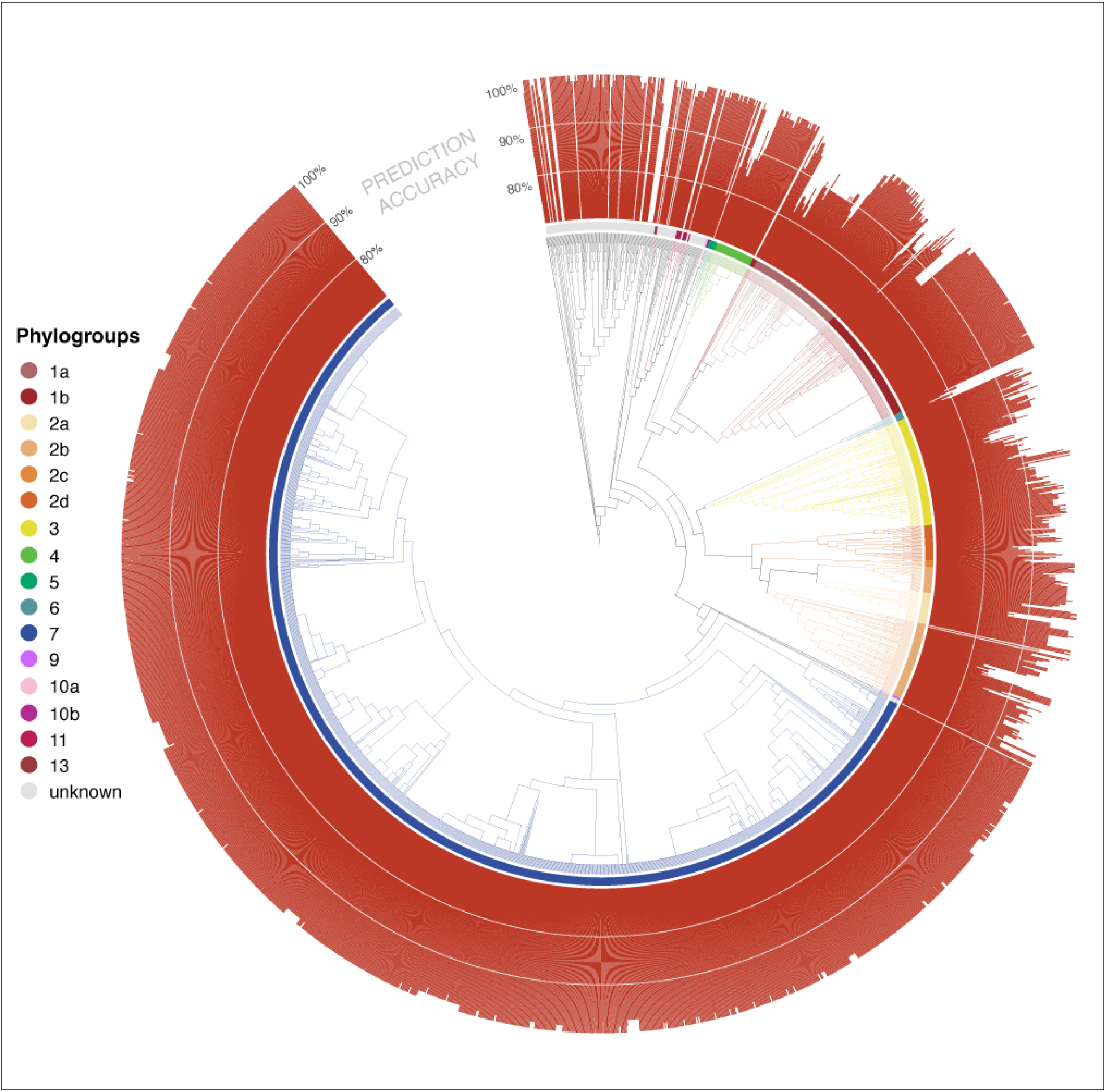
Similarity between individual type III effector repertoires and the consensus repertoire at the 98% ANI level. Height of the bar represents percent agreement, scaled from 70-100%. Genomes with no bar represent singletons at the 98% level, and thus no consensus repertoire could be calculated. Innermost ring colors designate phylogroups, as seen in fig. 1.

To further test the feasibility of T3E repertoire prediction, five genomes not included in our initial training set underwent in-silico PCR for the nine primers tested above, classified using the trained naïve Bayes classifiers, and screened for T3Es. Actual T3E repertoires were then compared to the average repertoire within the predicted LIN group of the unknown strain, and prediction accuracy for the repertoire was assessed (Fig. 3). Classification resulted in the placement of three strains into phylogroup 3, one in phylogroup 4, and the last into a cluster containing strains from both phylogroup 2c and 2d. For isolates placed into phylogroup 3, some primer sets were only able to classify to 97%, while other classified to 98% ANI, allowing us to also compare the accuracy of gene content predictions at difference classification resolutions. Prediction accuracy, measured as the proportion of effector protein subfamilies in the unknown isolate that could successfully be predicted by their frequency within the predicted cluster, ranged from 79.2-94.8% (fig. 4). Interestingly, increasing classification resolution had a small but negative effect on gene prediction accuracy in all three instances investigated. We do not know how to explain this observation. It could be a result of reduced sample size that is inherent in increasing resolution that introduces noise and decreases the prediction accuracy, further studies will be necessary to confirm and understand this pattern. Regardless, this result suggests that it might be beneficial to look at multiple classification levels when predicting gene content rather than only making inferences from the highest classification level possible. This result also suggests that the majority of primers tested here provide sufficient resolution for functional prediction purposes.

**Figure 4.**
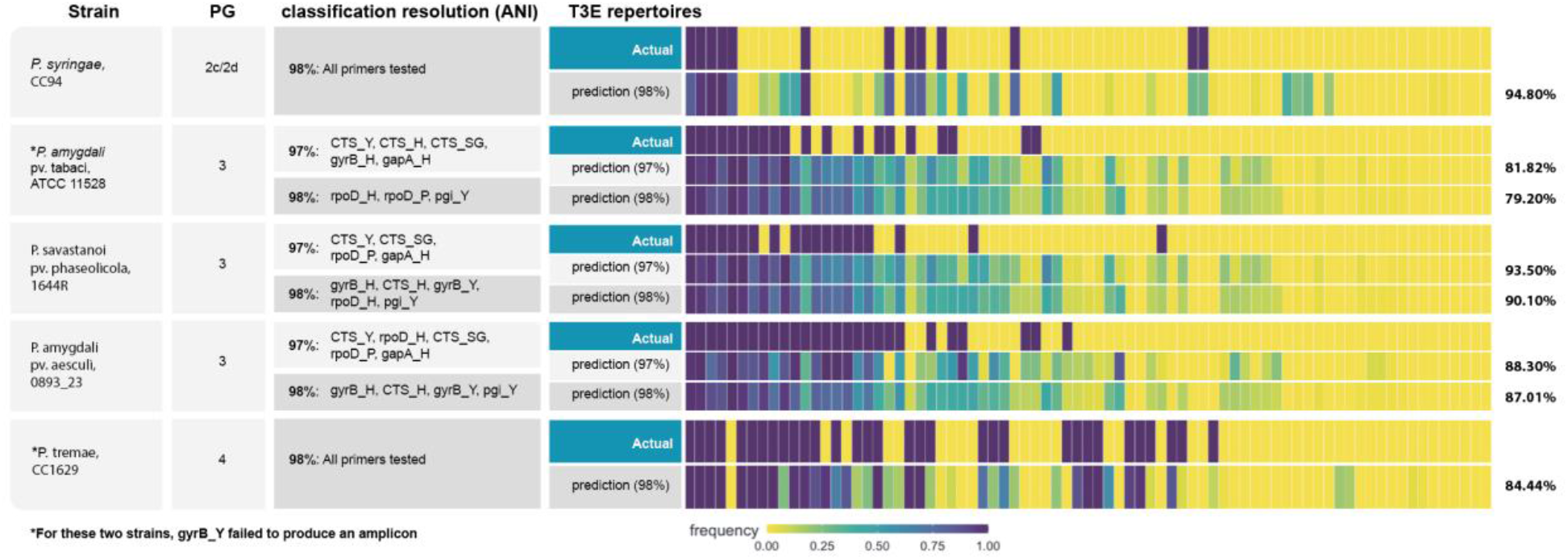
Classification based on single marker genes allows for accurate prediction of type III effector repertoires. The third column shows classification resolution obtain by each primer set. Heatmaps show the actual repertoire, with purple indicating presence and yellow absence of the gene subfamily, along with the average repertoire displayed by genomes within the genomic cluster that the unknown isolate was place into. Percentage of effector proteins for which the prediction agrees with the actual repertoire of the unknown isolate is shown to the right.

As more PSSC genomes are sequenced and deposited in public repositories, it is likely that our ability to predict T3E repertoires, as well as the presence of other important virulence factors, from amplicon sequencing will improve. This, coupled with recent research that indicates that host range can in part be inferred from virulence factors such as T3Es (Baltrus et al. 2012; Ferrante et al. 2009; Hulin et al. 2018) suggests that amplicon sequencing remains a powerful method for studying disease dynamics, predicting pathogen spread, and rapidly detecting problematic PSSC strains.

## CONCLUSION

In this study we set out to compare PCR primer sets designed to amplify broadly within PSSC. We found that there were significant differences in amplification rates that raise questions about the utility of some commonly used primers. However, we also found that classification resolution was relatively consistent between the primers tested (classification at the 97-98% ANI level).

The high resolution obtained from our classification models led us to investigate the potential of single amplicon sequences for prediction of type 3 effector protein subfamilies. we showed that with 79-94% accuracy, we were able to correctly predict effector repertoires of five PSSC strains. These results highlight the importance of continued isolation and sequencing of plant pathogens as a source of data to be leveraged in the future for more efficient and informative screening assays. Based on our findings here, we currently recommend the primer sets gapA-H, gyrB-H, and PGI-Y for single amplicon sequence typing of isolated PSSC strains.

## ACKNOWLEDGEMENTS

This material is based upon work supported by the National Institute of Food and Agriculture, U.S. Department of Agriculture, through the Northeast Sustainable Agriculture Research and Education program under subaward number GNE20-232.

CF is supported by the College of Agricultural Sciences (Penn State) and the department of Plant Pathology and Environmental Microbiology (Penn State) through the mBiome initiative.

EC is supported by Hatch fund 4710 entitled Fates of Soil Carbon and Nitrogen in Agricultural Systems in the College of Agricultural Sciences (Penn State) and the Huck Institute for the Life Sciences (Penn State). Authors also acknowledge the Penn State Microbiome Center, a community of scholars and students who coordinate and accelerate interdisciplinary discovery and applications to establish long-lasting resources in the field of microbiome research.

Additional support for K.L.H. came from the USDA National Institute of Food and Agriculture and Federal Hatch Appropriations PEN04648 (accession no. 1016871) and start-up funds through The Huck Institutes for the Life Sciences and the College of Agricultural Sciences at Penn State.

## DATA AVAILABILITY STATEMENT

T3E sequences detected by HMMER and used to assess repertoire content are available at https://doi.org/10.5281/zenodo.7249289. In-silico PCR results for primer sets exhibiting >50% amplification rate are available at https://doi.org/10.5281/zenodo.7249269. All other data is available upon request.

## SUPPLEMENTAL FILES

*Supplemental table 1* Data associated with genomes used in this study. Includes assigned phylogroups, ANI-based cluster assignments, and accession numbers and metadata extracted from GenBank.

*Supplemental table 2* Amplification rate for each primer set tested, by phylogroup

